# Independent Markov Decomposition: Towards modeling kinetics of biomolecular complexes

**DOI:** 10.1101/2021.03.24.436806

**Authors:** Tim Hempel, Mauricio J. del Razo, Christopher T. Lee, Bryn C. Taylor, Rommie E. Amaro, Frank Noé

## Abstract

In order to advance the mission of *in silico* cell biology, modeling the interactions of large and complex biological systems becomes increasingly relevant. The combination of molecular dynamics (MD) and Markov state models (MSMs) have enabled the construction of simplified models of molecular kinetics on long timescales. Despite its success, this approach is inherently limited by the size of the molecular system. With increasing size of macromolecular complexes, the number of independent or weakly coupled subsystems increases, and the number of global system states increase exponentially, making the sampling of all distinct global states unfeasible. In this work, we present a technique called Independent Markov Decomposition (IMD) that leverages weak coupling between subsystems in order to compute a global kinetic model without requiring to sample all combinatorial states of subsystems. We give a theoretical basis for IMD and propose an approach for finding and validating such a decomposition. Using empirical few-state MSMs of ion channel models that are well established in electrophysiology, we demonstrate that IMD can reproduce experimental conductance measurements with a major reduction in sampling compared with a standard MSM approach. We further show how to find the optimal partition of all-atom protein simulations into weakly coupled subunits.

**Significance Statement:** Molecular simulations of proteins are often interpreted using Markov state models (MSMs), in which each protein configuration is assigned to a global state. As we explore larger and more complex biological systems, the size of this global state space will face a combinatorial explosion, rendering it impossible to gather sufficient sampling data. In this work, we introduce an approach to decompose a system of interest into separable subsystems. We show that MSMs built for each subsystem can be later coupled to reproduce the behaviors of the global system. To aid in the choice of decomposition we also describe a score to quantify its goodness. This decomposition strategy has the promise to enable robust modeling of complex biomolecular systems.

---

The dynamics of proteins and their functions are of key importance for biology. Molecular dynamics (MD) simulations are a popular method for interrogating the motions of proteins in various environments. A well-known limitation of MD is the timescale mismatch between simulations and real life. Despite advances in computer hardware and algorithms, extreme timescale simulations remain orders of magnitude shorter than many relevant protein processes. Since one requires sufficient numbers of observations in order to obtain statistical confidence, various strategies have been developed to address this. One approach, building Markov state models (MSM), enables the construction of simple models of long-timescale molecular kinetics from many short off-equilibrium MD simulations (1–6) – see Refs. (7, 8) for thorough reviews. MSMs have successfully been built to obtain compact and yet accurate representations of the kinetics of full proteins (9–16), protein-ligand (17–22) and even protein-protein systems (23).

Although MSMs have significantly helped to reduce the MD sampling problem, the fundamental problem that arises from modeling increasingly large biomolecular systems remains. As protein complexes become larger, the number of uncoupled or weakly coupled subsystems increases. If each of these subsystems contain two or more substates, the number of global systems states increases exponentially (24). Therefore, any model treating the whole system by means of a global state poses requirements on the MD sampling that are fundamentally unscalable. This poses an inevitable problem as evolution tends to lead to increased biological complexity, including the optimization of processes through the formation of protein complexes and puncta (25–28).

In practice, many current models based on MD simulation of large biomolecular systems take the pragmatic approach of ignoring most of the system’s dynamics. For example, if one is interested in how an ion channel conducts ions across a membrane, it may be sufficient to prepare the system in a state of interest and collect sufficient statistics of ion passages and perhaps local confirmational changes of the selectivity filter residues, rather than trying to sample global conformational rearrangements of the protein complex on much longer timescales. However, our field has a collective interest in developing whole cell and systems modeling for *in silico* medicine which will necessitate the eventual understanding of these large systems in a way that characterizes how all their components interact, undergo transitions, and can be influenced by e.g., drug molecules, phosphorylation, and/or glycosylation states.

To this end, Noé & Olsson (24) have recently proposed dynamic graphical models which attempts to decompose protein systems in a way similar to Ising or Potts models – subsystems with states or “spins” that are coupled to one another. Dibak et al (29, 30) have developed a coupling of MSMs with reaction-diffusion dynamics in order to establish an infrastructure in which MSMs can be integrated into whole-cell models. Here we ask a more fundamental question, the answer of which is important to all these integrative approaches: given a large biomolecular system, how should we decompose it into subsystems, such that these subsystems can be described by independent or weakly coupled MSMs?

Fragmenting proteins at the modeling stage is compatible with prior experience as macromolecules are often sub-divided into structural or functional subunits (31). There is also evidence that proteins are decomposable into “quasi-independent groups of [spatially adjacent] amino acids” coined “protein sectors” (32). Furthermore, experimental studies on drug binding or protein functional characterization often use isolated domains or monomers with great success (33).

Estimating an MSM on the decomposed protein can significantly reduce the total sampling necessary. From concepts in statistical physics, given a polymer of length *N* where each subunit exists in one of *k* states, the total conformational space is expressed as *k*^*N*^ (cf. Fig. 1). Modeling subsystems of a constant size effectively restricts the number of states that need to be sampled reversibly to a constant. Therefore exponentially less sampling is required for modeling smaller subsystems as compared to a global model (15, 24).

**Fig 1.**
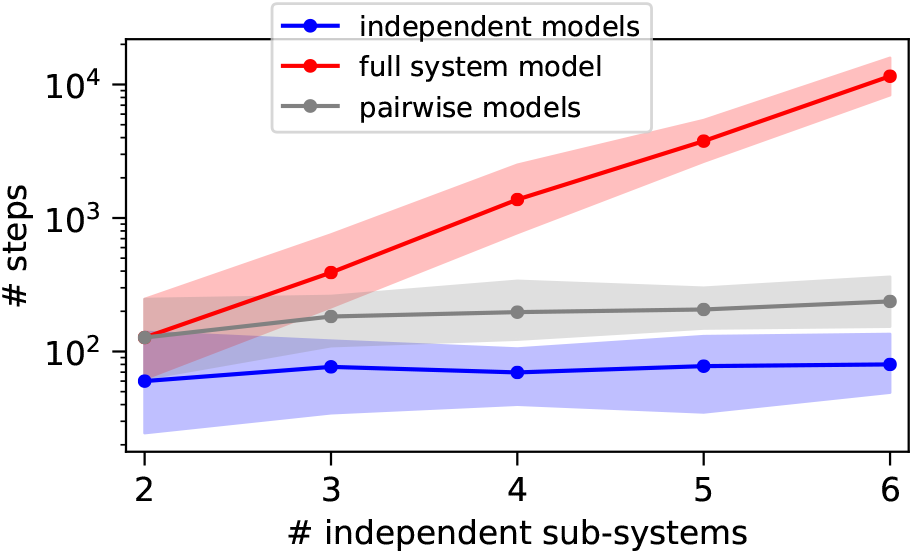
Scaling behavior of toy system consisting of *n* independent subsystems with 3 states each (SI Appendix, Toy models). Number of steps required to reversibly sample all transitions shown for proposed independent models (blue line), full system model (blue line) and pairwise models that are needed for computing the *dependency* score (gray line). Shadowed areas indicate 95% confidence intervals.

In this paper, we develop a mathematical framework of decomposing MSMs into local subsystem MSMs, termed independent Markov decomposition (IMD, Sec. IMD), and propose a measure of decomposition quality, the *dependency* score (Sec. MSM score of independence). We speculate that the IMD strategy can forge a new connection to other uses of MSMs such as those employed by the neuronal and cardiac modeling communities. There, phenomenological MSMs parameterized from electrophysiology data are used to predict the behavior of action potentials (34–39). In Sec. Tetrameric ion channel we describe how a decomposed MSM can be connected to a phenomenological MSM. This new connection between fields brings us closer to our goals of understanding these large systems and their behaviors, advancing *in silico* medicine. We further showcase how the *dependency* score can be used to find an optimal partition of a system that does not come with clearly defined independent subunits (Sec. Ion channel parti- tion). We validate our approach with a toy model, showing that the decomposition approximation is high quality and that the proposed validation score works even with limited data (SI Appendix, Toy models). Finally, we demonstrate its applicability to an all-atom MD dataset of the Synaptotagmin-C2A domain (Sec. Synaptotagmin-C2A partition) and derive the graph structure of inter-residue *dependencies*.

## Independent Markov Decomposition

We first describe IMD for discrete-state MSMs before generalizing it to time series with continuous descriptors.

### Markov State models

An MSM consists of a discretization of molecular state space into a disjoint set of states {*S*_1_, …, *S*_*n*_} and a Markov chain transition matrix ***P*** (*τ*) modeling a memoryless jump process between these states. We can express whether we are in the *i*th state or not by using indicator functions:

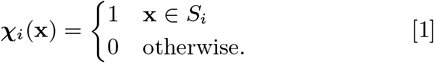

The vector **𝒳** = [**𝒳**_1_, …, **𝒳**_*n*_]^┬^ is thus a “one-hot encoding” that maps the continuous state **x** to the MSM discretization. For this or any other choice of features **𝒳** we can compute the instantaneous and time-lagged correlation matrices ***C***_00_ = **∑**_**t**_ **𝒳** (**x**_*t*_) **𝒳**^┬^(**x**_*t*_) and ***C***_0*τ*_ =**∑**_**t**_ **𝒳** (**x**_*t*_) **𝒳**^┬^(**x**_*t*+*τ*_), respectively. For a fixed state discretization, the transition matrix that has maximum likelihood and also maximizes the variational approach of conformation dynamics (VAC) (40) is:

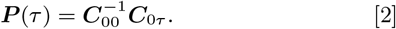

Let **p**_*t*_ denote the probability distribution of being in any of the *n* states at time *t*, for example **p**_0_ = [1, 0, …, 0] denotes that we always start in state 0 at time 0. This vector can be evolved in time using the transition matrix, until it converges to the equilibrium distribution **π** = lim_*t*→∞_ **p**_*t*_:

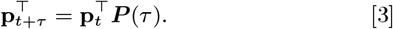

An important concept for optimizing the parameters or hyperparameters of MSMs and other Markovian kinetic models is the variational approach for Markov processes (VAMP) (41). VAMP finds that a Markovian model that best approximates the high-dimensional continuous dynamics maximizes the VAMP-*n* score:

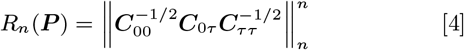

where we can either use *n* = 1 for the trace norm or *n* = 2 for the Frobenius norm. If we run molecular dynamics at equilibrium conditions, and we can employ correlation matrix estimators that provide ***C***_00_ = ***C***_*ττ*_ and symmetric ***C***_0*τ*_ (detailed balance). In this special case, VAMP becomes the VAC mentioned above, and the variational score simply becomes 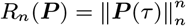. In other words, the optimal MSM is the one that maximizes the trace or the Frobenius norm of the transition matrix, which is equivalent to maximizing its eigenvalues. Since the eigenvalues equal the normalized time-autocorrelation of the slowest processes (1, 42), the VAC tries to find the Markovian model that best resolves the slowest processes of the molecular process under investigation (40, 43). For a fixed state space discretization, optimizing the VAC results in the MSM estimator (2). If we also want to search over different state space discretizations, we can use VAC or VAMP as a score in a hyperparameter optimization problem (44) or optimize the VAMP score while representing **𝒳** with deep neural networks, leading us to VAMPnets (45).

### Independent Markov decomposition

Now we move beyond the common concept of modeling the dynamics of the entire molecular system by a single MSM and instead try to decompose the system into almost independent MSMs. Let us start with the simple example shown in Fig. 2a, where a molecule consists of two domains *A* and *B* that are each described by a two-state MSM describing whether the domain is “closed” (*α, β* = 0°) or “open” (*α, β* = 90°). We assume that the kinetics of both domains are statistically independent, i.e. each domain switches states independent of the states of the other one – we simultaneously have ***p***_*A,t*+*τ*_ = ***P***_*A*_(*τ*)***p***_*A,t*_ and ***p***_*B,t*+*τ*_ = ***P***_*B*_ (*τ*)***p***_*B,t*_ (Fig. 2b). As the MSMs *A* and *B* are statistically independent, the probability distribution of the entire system follows Eq. (3) with

**Fig 2.**
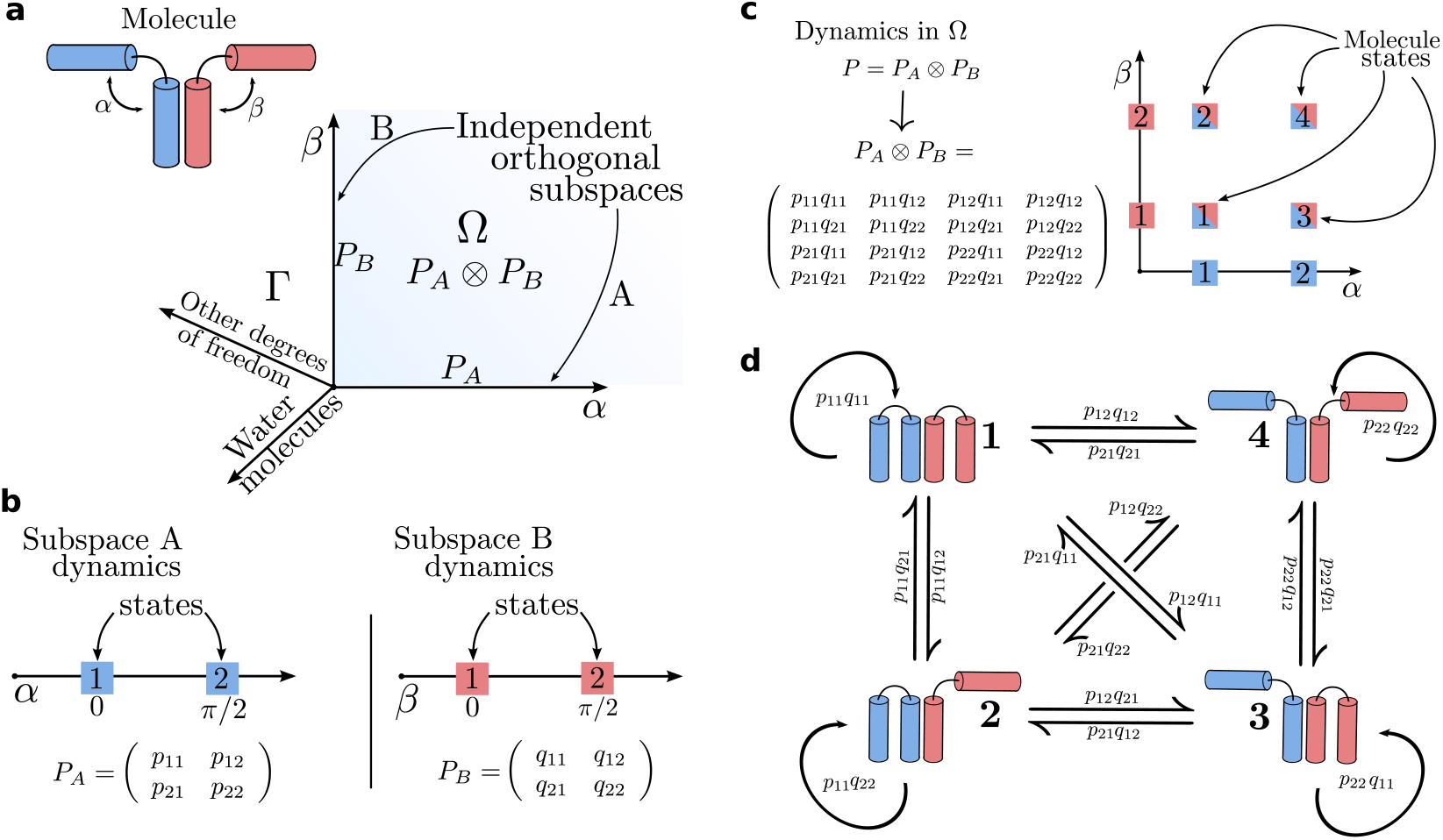
Operator decomposition and discretization on a test molecule. a) A test molecule is decomposed into two subsystems (blue and red). The two angles *α* and *β* span subspaces *A* and *B* corresponding to the two subsystems, respectively. The space Γ is composed of all system degrees of freedom. The space Ω is the Cartesian product of *A* and *B* and its dynamics are described by Perron-Frobenius operators *P*_*A*_ and *P*_*B*_, respectively. The dynamics in Ω are given as the tensor product *P*_*A*_ ⊗ *P*_*B*_. b) The molecule has metastable states at *α* = 0, *π/*2 and *β* = 0, *π/*2; the subspaces *A* and *B* can be discretized into MSMs with transition probability matrices ***P***_*A*_ and ***P***_*B*_. The quantities *pij* and *qij* are the transition probabilities from state *i* to *j* of subspaces *A* and *B*, respectively. c) The discretized dynamics in Ω are given by the tensor product ***P***_*A*_ ⊗ ***P***_*B*_, yielding the four states of the full molecule. d) Illustration of the four possible states of the molecule and the transitions between them.

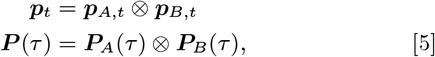

where ⊗ is the Kronecker product (46) (see SI Appendix, Markov operators). The vector ***p***_*t*_ now contains the probabilities of being in the four combinatorial states (*A* and *B* open, *A* open and *B* closed, *A* closed and *B* open, *A* and *B* closed), and ***P*** (*τ*) is the 4 × 4 transition matrix between these combinatorial states whose transition probabilities are simply products of the individual transition events in subsystems *A* and *B* (Fig. 2c, d).

The power of this approach is apparent when comparing Figures 2b and c: If the dynamics in *A* and *B* are independent or almost independent, we can estimate the sixteen transition probabilities that parametrize the whole system using only the eight elements of the transition matrices of the subspaces. This advantage increases exponentially in larger systems: If we have *N* (almost) independent domains with *m* states each, distinguishing all states would require us to sample and estimate an exponential number of order of *m*^2*N*^ transitions, while a decomposition into independent MSMs reduces this to a polynomial number of *Nm*^2^ transitions which can be scaled to large systems.

The above example trivially generalizes to *N* systems with 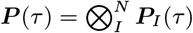. We note that it is customary to dismiss variables of the full state space Γ (Fig. 2a) that are assumed to average quickly, e.g., solvent degrees of freedom. Thus the modeled space Ω in practice only encompasses the variables of interest, e.g., internal coordinates of a protein system.

### An MSM score of independence

In practice, subdomains of biomolecules or biomolecular complexes will not be exactly independent. Moreover, the identification of a domain decomposition into almost independent subdomains is a non-trivial task. To enable algorithmic determination of almost independent subdomains, we develop an independence score which quantifies decomposition validity.

To this end we come back to the variational approach Eq. (4). Conveniently, matrix norms follow simple rules when applied to a Kronecker product (SI Appendix, Markov operators). In practice, we will apply the trace and Frobenius norms that correspond to the VAMP-1 and VAMP-2 scores of the Koopman operator. The VAMP-2 score has successfully been used in many practical applications (16, 45, 47, 48). If our molecular system consists of *N* independent subdomains such that its global MSM is a Kronecker product of *N* subspace MSMs as described above, its VAMP score is the simple product of VAMP scores (SI Appendix, VAMP score decomposition):

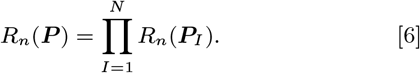

Here, *R*_*n*_(·) denotes the VAMP-*n* score of the transition operator. It could be the trace norm (VAMP-1) or Frobenius norm (VAMP-2) of the associated transition matrix. Note that Eq. (6) is a necessary but not a sufficient condition for Markov independence. Significant deviations from equality in Eq. 6 indicate that the assumption of independence is invalid. However, in particular if separate MSMs ***P***_*I*_ can probe the same molecular features it is possible to satisfy Eq. (6) even though the subsystem MSMs are not statistically independent. Eq. (6) must therefore always be used in conjunction with appropriate constraints. Here, we choose between different ways to assign independent molecular features to different MSMs and check which of these assignments best satisfies Eq. (6).

In practice, we want to perform an IMD because often we can not compute the global MSM ***P*** due to limited sampling (Fig. 1), and we consequently do not know *R*_*n*_(***P***). Therefore, we choose to only check the equality of Eq. (6) on pairs of subsystems *A, B*, i.e., *R*_*n*_(***P***_*A,B*_) = *R*_*n*_(***P***_*A*_)·*R*_*n*_(***P***_*B*_). We then search over possible partitions of the molecular system into subsystems by evaluating the graph of pairwise *dependencies d*(*A, B*):

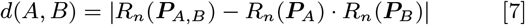

In practice, computing ***P***_*A,B*_ involves a new estimate of the transition probability matrix in the joint space of two systems. We show that our measure scales well with respect to limited sampling (also compare SI Appendix, Toy models).

The product in Eqs. (6) and (7) is purely a result of the chosen basis set of MSMs (Eq. (1), SI Appendix, VAMP score decomposition). In practical situations, it is desireable to find a decomposition directly based on molecular features such as distances or contacts instead of carrying out an MSM discretization and estimation for each subsystem. When considering more general features **𝒳**, there are two main changes to discrete-state MSMs: (i) observables are propagated by a different operator, called Koopman operator (49, 50), (ii) the joint space of observables is easiest described by “stacking” obserable feature vectors rather than by defining an MSM discretization on the combinatorial space. For example, if 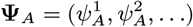 and 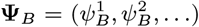 are the one-dimensional time series of features *ψ* ∈ ℝ of two systems *A* and *B*, the joint space would be spanned by 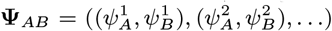. The transfer operator that describes the independent dynamics in joint space is thus a block matrix of its constituting independent sub-operators (also called a direct sum, see SI Appendix, Markov Operators for details). This also means that independent subsystem features are not correlated. Please note that “stacking” in the MSM formulation would produce probability vectors not normalized to 1 and yield invalid (i.e. not irreducible) MSM transition matrices in the joint space.

The trace and Frobenius norm of the Koopman operator thus decompose as sums such that the dependency score reads

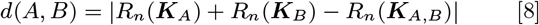

where ***K***, the Koopman operator, takes the place of the transition matrix ***P***. See SI Appendix, VAMP score decomposition for the derivation. Please also note that working in observable space entirely rules out discretization artifacts.

## Results

### Modeling a tetrameric ion channel using IMD

In cardiac electrophysiology, Markov models have been used to model phenomenological data from ion channels (37–39). Ion channels are transmembrane proteins that respond to physiological stimuli and selectively control the flow of ions in excitable cells. Upon a change in membrane potential, voltage-gated ion channels undergo conformational changes which modulate ionic conductance. The symphony of ion channels collectively facilitate the propagation of electrical signals in excitable tissues, such as the heart and brain, and are important drug targets (51, 52).

The plethora of both experimental measurements of ion channel properties sets the stage for computational simulations to provide molecular details and mechanistic insights (53). While it is possible to fit a phenomenological MSM using data from electrophysiological experiments, atomistic modeling remains out of reach due to the long timescale of channel opening. This is because single gate activation events are rare, and many ion channels have multiple gates which need to activate concurrently. Reversible sampling will further be hampered by a combinatorial number of pathways that lead to a fully open channel. We propose that for cases of non-cooperative gates, IMD can help solve this problem, which we demonstrate in the following series of numerical experiments.

We consider a voltage-gated tetrameric potassium ion channel with four identical subunits, each with a voltage sensor. To construct an IMD, we exploit the independence of individual subunits or gates and partition accordingly (Fig. 3a1). This produces four matrices ***P***_*i*_ ∈ ℝ^2*×*2^, 1 ≤ *i* ≤ 4 that describe individual gate opening and closing. As derived above, the Kronecker product of subsystem transition matrices yields a transition matrix ***P*** ∈ ℝ^16*×*16^ of the full ion channel (Fig. 3a2). The 16 states enumerate all possible combinations of open and closed gates of the full ion channel, a state space referred to as 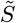 in the following. We note that this decomposition is only possible between non-cooperative domains.

**Fig 3.**
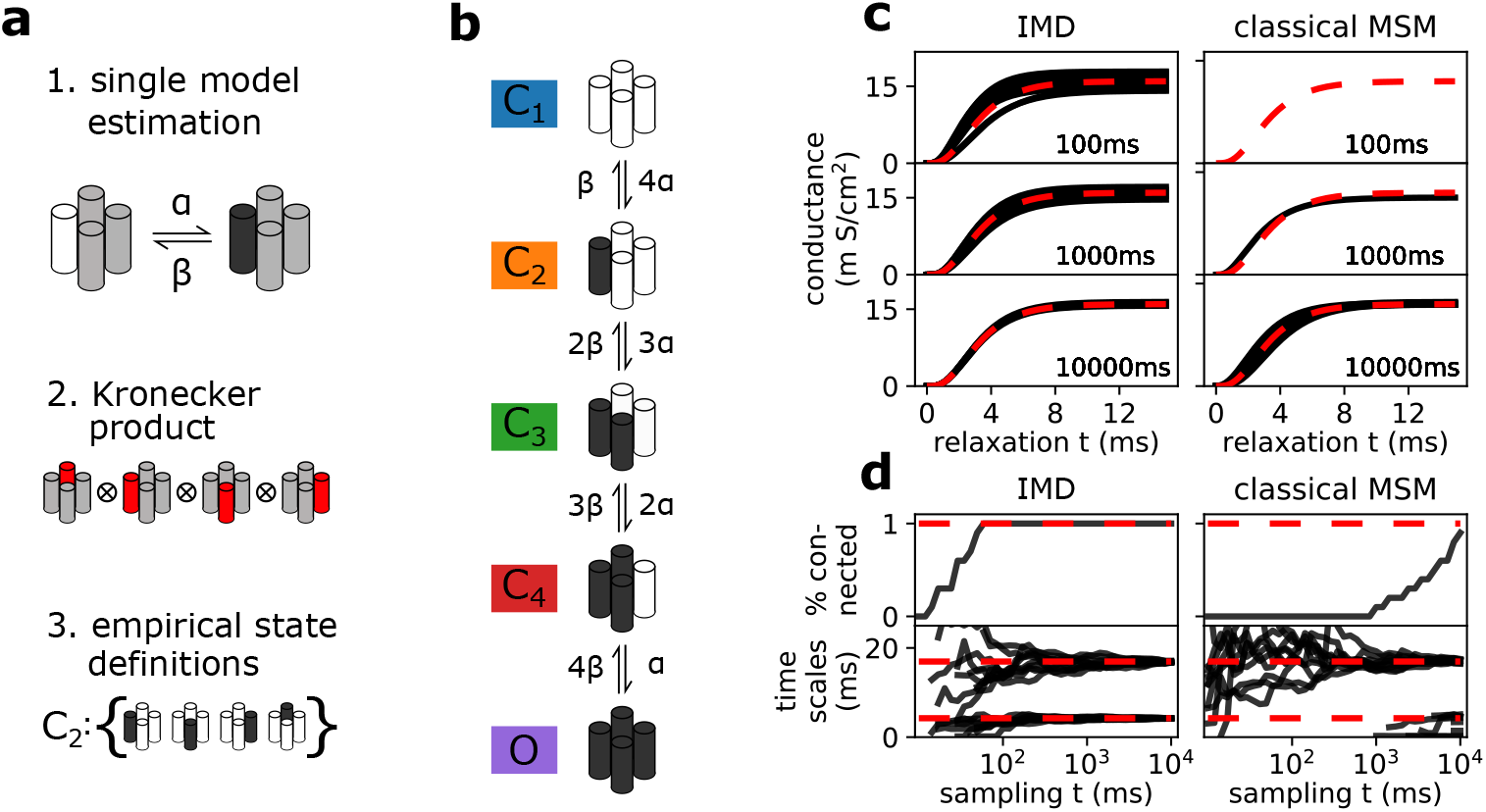
Reconstructing the Hodgkin-Huxley model from a simple discrete model. a) Pipeline of steps required to assemble a full channel model from a single subunit model that describes opening and closing of a single subunit in the vicinity of the others (step 1). Kronecker product between all four sub-unit models assembles a model that still distinguishes between all combinatorial states (step 2). Empirical state definitions account for channel symmetries (step 3). Black denotes open, white closed, and gray undefined subunit. b) Graphical depiction of full channel model in empirical state space. Note the symmetry of the channel, i.e. that at this stage only the number of open subsystems is known. c) Relaxation from a closed state into the native state at 63 mV. We show conductance predicted by IMD (left column) and classical MSM (right column) using different amounts of sampling. Note that the classical approach only yields results in the high sampling regime where all empirical states are connected. Results are compared to the original Hodgkin-Huxley model (red dashed line). d) Sampling time necessary to estimate a decomposed MSM (left column) compared to a classical full system MSM (right column) for ten realizations of the Markov chain. We show the percentage of fully connected models in our ensemble of realizations (top row) and the 1st and 4th implied timescale computed from it (bottom row). Note that for the classical MSM, extreme amounts of sampling are necessary to even estimate all system-inherent implied timescales.

We construct a mapping to assign the 16 states of the transition matrix ***P*** to those of a phenomenological MSM. Our reference empirical model is the one developed in Ref. (54) for this channel (Fig. 3b). In Ref. (54), channel symmetry is used to define the full system states accordingly:

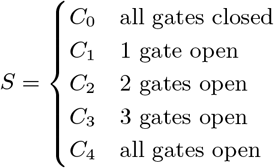

Mapping of the transition matrix into the space of these empirical states can be obtained by converting the empirical state definitions into crisp membership vectors **𝒳**_*s*_ ∈ {0, 1}^5^, with each element indicating which empirical configuration a full system configuration 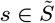 belongs to. For example, the membership vector describing any state *s*_*k*_ with one open gate would be 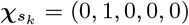, i.e., these states are associated to macroconfiguration *C*_1_. The full membership matrix is constructed by stacking 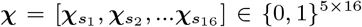. Subsequently, the transition-matrix is coarse-grained following (55, 56) 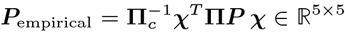 with **Π** = diag(**π**) the diagonal matrix of the stationary distribution *π* in full space and in empirical space **Π**_*c*_ = diag(**𝒳**^*T*^ π).

Choosing rates *α* and *β* from the original work by Hodgkin-Huxley (34) at a voltage of 63 mV, we produce a simple discrete model. Using this model, we can generate sample trajectories from which to construct MSMs in accordance with Sec. Computational experiments. We estimate a model for the full system from this data by applying the aforementioned pipeline. Using this derived full system model, experimental observables from electrophysiology experiments can be assessed by relaxation of the Markov chain from a non-equilibrium distribution (e.g. a closed configuration) into the equilibrium at this particular voltage (57, 58). We start from a configuration of fully closed states and further assume that the channel only conducts ions if it is open, i.e., our observable is only non-zero for the open state. This experiment is the computational analog to a voltage jump experiment from resting to +63mV in voltage clamp mode. Shown in Fig. 3c, the modeled conductance of the channel over time is reported. The predicted conductance time-series is compared with the numerically integrated ordinary differential equation for the potassium ion channel derived by Hodgkin and Huxley (34). We find that IMD can accurately reproduce the full channel dynamics.

IMD models were built by separately fitting four single gate trajectories (i.e. a full system trajectory split into its subsystems) and assembled using the aforementioned steps. For comparison, traditional MSMs were fit to sample trajectories computed from the full system transition matrix in its empirical state definition. We note that we compare IMD sampling to the empirical 5-state formulation (which does not resolve all 16 combinatorial states). In this way, we can rule out that the described sampling advantages of IMD are an artifact of exploited channel symmetry.

The reduction in the amount of sampling needed due to the use of IMD can be quantified in terms of the length of simulation required to form a fully connected transition matrix. In Fig. 3d we present the percentage of connected IMD estimated on an ensemble of ten realizations of the Markov chain and compare to a classical MSM. It is computed as a function of simulated time (in ms), i.e., shows how probable a modeler can estimate a connected Markov model, IMD or classical, from a fixed amount of sampling. We note that the classical MSM approach can only estimate all system-inherent implied timescales when all empirical states are reversibly sampled, i.e., only for very large amounts of data. The higher computational efficiency of IMD is evident from the much faster convergence of implied timescales as a function of simulation length (Fig. 3d), showing a reduction in sampling by three orders of magnitude, from tens of seconds to tens of milliseconds (Fig. 3d). For example, ionic conductance is reasonably approximated with 100 ms of sampling and the IMD approach (3c).

Here, we have presented an example where each gate operates independently. In practice, the gating behaviors of most ion channels are not completely independent, but are instead coupled. In this case, the decomposition yields an approximate model of the real dynamics, see SI Appendix, Weakly coupled systems for a discussion. The theoretical limit is posed by the assumption of stationarity that underlies MSM estimation. It is violated if external influences are strong and on similar timescales as the processes to be modeled. External influences that are much faster than the local dynamics are incorporated as an average over Markov states, similar to water molecules in regular MSMs. As demonstrated in the SI Appendix, Fig. S1, modeling of weakly coupled systems is possible in a robust fashion.

### Finding Independent Markov Partition for tetrameric ion channels

For our previous example, we prescribed a convenient partitioning scheme for the ion channel system. In contrast, in real-world situations a complex system may involve multiple independent subsystems but the coupling graph is unknown *a priori*. For instance, it might not be clear how to find independent protein segments of an unknown protein. A method is necessary to aid in the development of viable partitions which produce independent subsystems.

In this section we demonstrate how the *dependency* defined in Sec. MSM score of independence can be used as a score to bisect clusters of coupled subsystems from weakly coupled ones. The idea is to compute all possible pairwise *dependencies* between all subsystems and to use them as edge weights in a graph. If they exist, (almost) independent clusters of strongly coupled subsystems will be revealed by analyzing this graph. Once identified, these clusters might be modeled with single subsystem transition matrices within the IMD framework.

For the purposes of demonstration, we zoom out from a single channel protein to a membrane patch (Fig. 4a). In our setup, this patch contains a dimer of channels which we model to be coupled by a weak, cooperative coupling. Individual channels are modeled using the same parameters as the above ion channel model but contain the additional element of an external deactivation switch (Fig. 4b). In a cellular environment, such a switch could, for example, be an inhibitory ligand that binds and unbinds at a certain rate. It is modeled as a Markov process with probability 0.01 to change its state. The deactivation switch alters the conformational dynamics of each gate such that the probability to close or to stay closed is 95%. Thus, by construction, it is not possible to decompose a channel MSM into single gate MSMs because each gate is now coupled to the deactivation switch. Further, the strength of the intra-channel coupling can be controlled by a linear mixture parameter *λ*. The dynamics described above correspond to *λ* = 1, strong coupling. The coupling can be entirely deactivated by setting *λ* = 0. See SI Appendix, Dimer model for implementation details.

**Fig 4.**
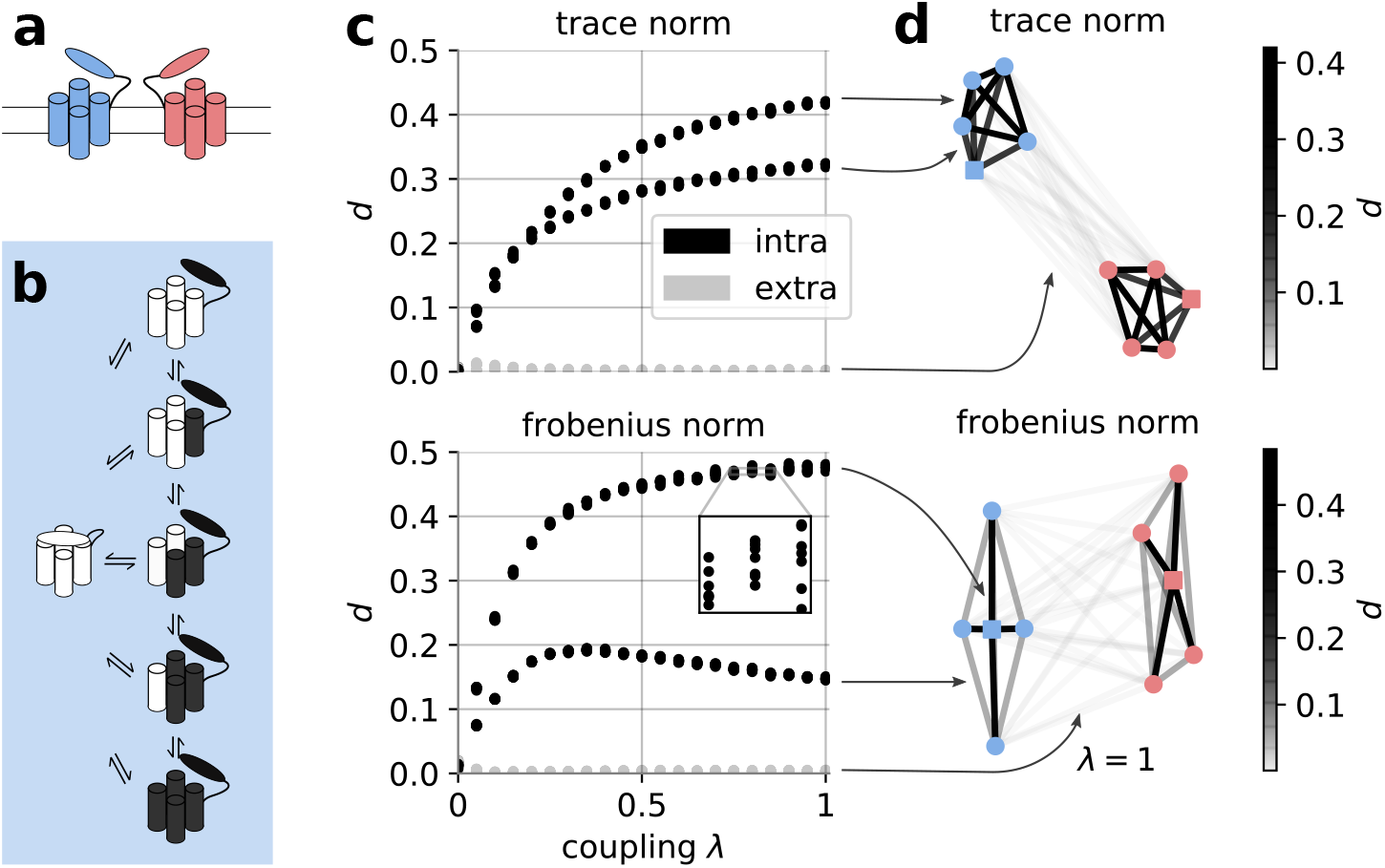
Visualization of channel dimer. a) Two channels located in a membrane. Each channel consists of four gates (akin to Hodgkin-Huxley model, depicted by cylinders) and one desensitization switch (depicted as appendix to four-bundle). b) States and possible transitions of individual channels (simplified, short lived switch-deactivated open states not shown). As both channels have the same dynamics, only one is shown as an example. c) Dependency score as a function of coupling strength as defined by linear mixture parameter *λ*. Color code: Grey denotes scores between two molecules, black intra-channel pairs. d) Graph of pairwise dependencies between all channel subunits for *λ* = 1. Edges are color coded according to *dependency* scores between two systems. Nodes belonging to a single channel are color-coded accordingly, squared nodes represent deactivation switches.

We generate discrete time series data from a transition matrix that models a dimer with these properties (SI Appendix, Dimer model and Computational experiments). From the data, the *dependency d* is computed for all possible pairs of subsystems. This involves the estimation of transition matrices for two isolated subsystems and comparing them with the transition matrix estimated in the joint space using Eq. 7. For example, one such pair could be the deactivation switch of one channel and a gate of the other channel.

A natural representation of these pairwise norms between subsystems is a graph. It is formed by nodes (subsystems) and *dependency*-weighted edges; no assumption about its structure is made (e.g., that it is a fully connected graph). For the numerical experiment described in this section, our analysis yields the graph shown in Fig. 4d.

The graph is visualized by positioning the subsystems or graph nodes with the Fruchterman-Reingold algorithm (59, 60) which is sensitive to the edge weights. This means that subsystems with high *dependency* are grouped together. This helps us to visually identify clusters of coupled subsystems. Groups of subsystems that are far apart in this representation are coupled relatively weakly. We find that *dependencies* between subsystems of the same channel are significantly larger than zero while inter-channel interactions yield *dependencies* close to zero (see Fig. 4d). Further, reducing the coupling strength within a channel does not alter our qualitative results (Fig. 4c). The observed bifurcation of *dependencies* is due to the two types of coupling in the system (gate-gate vs. gate-deactivation switch) and is a feature of the dimer model system.

In summary, our results show that we can learn the connectivity of a network of subsystems from discrete, simulated time series data. In particular, the *dependency* score provides an approach to find an optimal partition of a system with multiple types of coupling.

### Finding Independent Markov Partition for all-atom simulations of Synaptotagmin-C2A

To showcase the applicability of the *dependency* score, we apply our method to a 180 *µs* molecular dynamics data set of the C2A domain of Synaptotagmin-1 (Syt). Syt is a crucial player in the neurotransmitter release machinery (61). In our previous study we have found that single loops of its C2A domain can be described independently of each other using a hand crafted partition (15). Here, we attempt to find an optimal partition by using the *dependency* score at the residue resolution (Sec. Application to MD dataset). Instead of working with MSM transition probabilities, we directly work in protein feature space in order to omit discretization artifacts.

We find that indeed, Syt-C2A can be partitioned into defined subunits, or conformational switches, using a VAMP-2 based *dependency* score (Fig. 5). This partition contains the conformational switches defined in our last study (15): In particular, the C78 switch (Fig. 5, blue) emerges as an independent cluster in the Fruchterman-Rheingold projection, confirming our previous results. However, even though two conformational switches in the Calcium Binding Region (CBR), CBR-1 and 2 together (Fig. 5, green and red), have low dependency to the other protein residues, describing these loops independently is an approximation that is only partially backed by this current study. Similar results are obtained when using a VAMP-1 based *dependency* (SI Appendix, Fig. S2).

**Fig 5.**
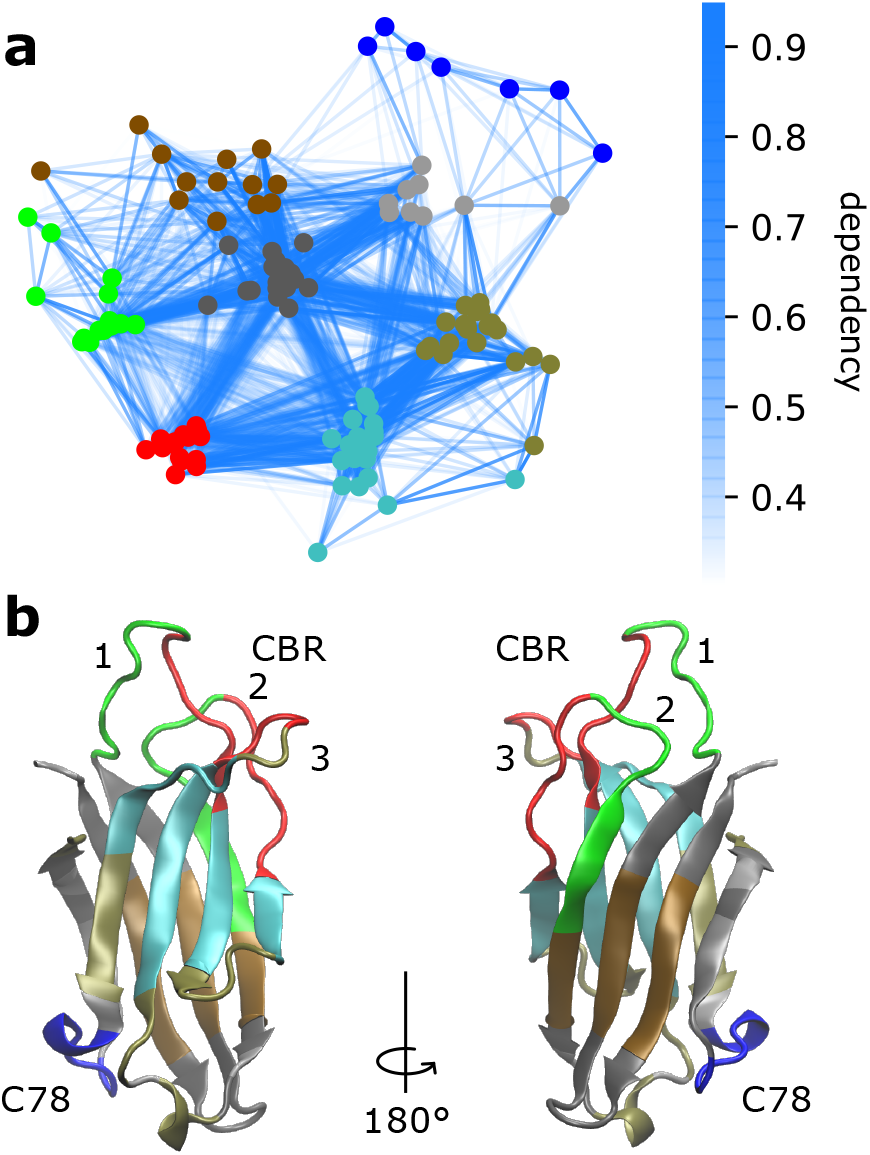
Dependency-network between residues of Syt-1 C2A depicted using a standard graph layout (Fruchterman-Rheingold algorithm). a) VAMP-2 normalized dependency network. Edge weights are indicated by colorbar. Nodes are colored according to an unsupervised classification by the *k*-means algorithm (*k* = 8). b) Visualization of protein structure with color coded segments from our VAMP-2 analysis (colors correspond to classification in panel a). VAMP-1 yields similar results (not shown here, see SI Appendix, Fig. S2).

## Discussion

Over the past several decades, MSM methodology has matured into a valuable tool for MD data analysis (1, 3, 4, 7, 8, 13, 20– 23, 42). For practitioners, modeling MD data with MSMs remains a non-trivial task, especially as researchers turn their focus towards the study of progressively larger biomolecular complexes. This development comes with an increasing number of (metastable) states that demands vast amounts of sampling time and hampers our attempts to rigorously model protein dynamics. We believe that the classical MSM method is reaching a point where the combinatorial explosion of states becomes a critical bottleneck. It is a fundamental problem that is inherent to any method which seeks to describe the global protein state (24).

One possible solution is to appreciate the notion of independent protein segments (32) and to split large systems into smaller, more manageable ones. In this spirit, we have proposed Independent Markov Decomposition. For practitioners, this means that, for example, an ion channel is modeled as a set of individual gates as opposed to a single protein. This approach approximates the system as a set of independent subsystems and is naturally agnostic to global system size. In this paper we have shown how the conceptual idea of decomposed MSMs relates to the underlying transfer operator formulation, what sampling advantages can be expected, and how to use the proposed *dependency* score to find an optimal partition of an unknown system.

Using the tetrameric potassium ion channel as a model system, we have shown that we can estimate a fully converged model with approximately three orders of magnitude less sampling when compared to a classical MSM. IMD therefore has the potential to leverage sampling efforts for large biological systems into a regime which is achievable with state-of-the-art simulation techniques and computer hardware. This effect is due to data being used more efficiently while small compromises are made by a mean-field-like approximation. For systems with potentially weak couplings, the validity of the approximation can be checked with our *dependency* score *a posteriori*. We further posit that due to the tremendous sampling advantages, the estimation errors introduced by weak couplings are likely to be smaller than the sampling error for classical global state MSMs.

Our results suggest that IMD improves the assessment of sampling convergence for large systems. As real-world MD datasets are usually very high dimensional, in practice, it is a non-trivial task to assess whether the sampling is converged. Often, researchers can only speculate by using semi-empirical tests, i.e., matching of high-level experimental observables to model predictions. IMD offers a more rigorous way to tackle this problem. For example, when modeling a single protein loop, it is much easier to see if the process is sampled reversibly, a question that can be difficult to answer with a classical MSM on global states.

Furthermore, we have proposed a *dependency* score that quantifies the coupling between two subsystems. As there is no general rule how to define protein subsystems, the *dependency* score serves as an objective function to judge IMD approximation quality and to find an optimal partition of unknown systems. In a numerical test system of a switched dimer model with weak cooperative coupling, the *dependency* score has robustly bisected clusters of strongly coupled subsystems from weakly coupled ones. It thus enabled IMD estimation without knowing the dependency graph structure *a priori*.

In order to optimally partition a system in practical applications, sufficiently large biomolecular system could be first partitioned into minimal subsystems such as residue sidechains. Scoring the dependency between these subsystems can reveal the structure of the dependency graph and thus give rise to a definition of (almost) independent protein segments. We have shown that for the C2A domain of Synaptotagmin-1, the *dependency* score can be used to identify clusters of subsystems that are linked relatively weakly between each other. These subsystems are similar to the conformational switches identified and independently modeled in Ref. (15). For future work, in particular for larger biomolecular complexes, it will however be desirable to incorporate experimental knowledge about size and properties of “protein sectors” (32).

An aspect excluded in this conceptual study is the discretization of MD data, a step which can be crucial in practical MSM applications (4, 62). We note that subsystem MSMs have smaller dimensionality and therefore discretization errors are smaller compared to the higher-dimensional full system. This implies that IMD may reduce discretization artifacts compared to classical MSMs. However, implications of the discretization error should be discussed when applying the *dependency* score as it is unclear how the error propagates to joint space probability estimates.

In this work, we propose that one way to keep pace with our interest in modeling large biological systems is by using a decomposition technique. For large systems, IMD is more data efficient and might be easier to apply than classical global state MSMs. We believe that interrogating local features, e.g., ligand binding pockets, instead of global system states can be more informative and give better predictions at reduced computational cost. Because this approach comes with all the established methods and software of the MD MSM community, we anticipate that IMD will have a broad application basis for *in-silico* biology.

## Materials and Methods

### Computational experiments

Gate opening and closing rates of the toy potassium ion channel were obtained from the Hodgkin-Huxley model. Under voltage clamp conditions and neglecting the sodium and leak currents, we are left with the potassium ion channel contribution. The current is given as follows,

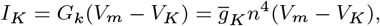

where *I*_*K*_ is the current, *G*_*K*_ is the conductance, 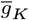 is the maximal conductance, *V*_*m*_ and *V*_*K*_ are the total transmembrane potential and potassium ion reversal potential respectively. Here *n* ∈ [0, 1] is a dimensionless quantity corresponding to channel activation. The time dependence of *n* is described using the following ODE,

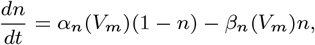

where *α*_*n*_ and *β*_*n*_ are the kinetic rates (*s*^−1^) of activation and deactivation respectively. In the original Hodgkin-Huxley model (34), the voltage sensitivity of the ion channel is modeled by the voltage dependence of the rates *α*_*n*_ and *β*_*n*_,

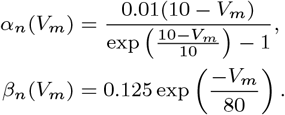

The term *n*^4^ is the joint probability that the four independent subunits of the tetrameric potassium ion channel are concomitantly open. Thus *α*_*n*_ and *β*_*n*_ are the kinetic rates for an individual subunit to open and close respectively. This set of ODEs were integrated using the odeint function provided by scipy (63) to serve as the ground truth for later comparison with IMD and MSM results.

We apply our framework to discrete time series data with known full system dynamics. For each system that we are using, details and generator matrix are given in the SI Appendix, Secs. Toy systems, Dimer model.

Generally, a transition matrix describing a (full) test system (possibly including couplings) is chosen, akin to ***P*** (*τ*) in Eq. 5. Time series are generated using the Markov chain sampler implemented in pyEMMA/msmtools (64). Subsequently, full system states are mapped to individual subsystem states, yielding subsystem trajectories which are parallel in time. Estimation of subsystem transition matrices (***P***_*i*_(*τ*) in Eq. 5) is followed by assembly of a full system transition matrix. The latter is utilized to extract full system observables such as implied timescales.

### Application to MD dataset

The protocol that was used to obtain MD simulation data and featurization of Syt-C2A is described in detail in Ref. (15). In particular, as in the cited study, we use heavy atom coordinates of the superposed protein. We are aware that this could potentially yield spurious correlations, however a) no better descriptor of the slow dynamics could be found and b) we want to ensure compatibility to our previous study.

Each residue is encoded as a vector of flattened coordinates *Y*_*i*_ and the *dependency* is computed on each pair of residues. The pairwise features are the stacked vectors [*Y*_*i*_, *Y*_*j*_]. Note that when directly working on coordinate features, unlike in the MSM examples, the *dependency* decomposes as a sum, not as a product (SI Appendix, VAMP score decomposition). Furthermore, the dependency is normalized to untangle the amount of kinetic variance from actual dependency, i.e.

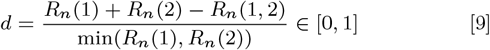

with *R*_*n*_(*x*) being the VAMP-*n*-score of residue *x*. The VAMP-*n*- score is computed with PyEMMA (64).

Note that in the case of high *dependency* scores, the two observable features might be proxies of the same process, however one of them could encode an additional one. Dividing by the min ensures we are only normalizing to the processes contained in both subsystem vectors.

## Data availability

The code that implements our discrete models and reproduces the presented results can be found in our GitHub repository. The molecular dynamics data set of Synaptotagmin C2A is available upon request.

## ACKNOWLEDGMENTS

TH thanks Moritz Hoffmann and Andreas Mardt (FU Berlin) for fruitful discussions. We acknowledge funding from Deutsche Forschungsgemeinschaft (SFB/TRR 186, Project A12; SFB1114, Project A04), the Berlin Mathematics center MATH+ (AA1-6), the Bundesministerium für Bildung und Forschung BMBF (BIFOLD), and the European Commission (ERC CoG 772230 “ScaleCell”). BCT and REA acknowledge support from the National Biomedical Computation Resource via NIH Grant P41-GM103426. CTL acknowledges support from a Hartwell Foundation Postdoctoral Fellowship. REA acknowledges funding from NIH R01 GM132826.

## Notes

Conflict of interest statement: The authors declare no conflict of interest.

### Competing Interest Statement

The authors have declared no competing interest.

