## Supplementary material for "Independent Markov Decomposition: Towards modeling kinetics of biomolecular complexes": SI appendix


#### Markov operators

##### Infinitesimal generator

Given a stochastic dynamical system, such as an MD simulation, the operator $P$ can be understood as a propagator of the probability density $f(x, t)$, where $x$ is on the phase space of the dynamical system. We can define $P$ using the infinitesimal generator $L$ [1]

$$\partial_t f = Lf. \tag{1}$$

This equation is a generalization to non-deterministic systems of the Liouville equation of statistical mechanics, which describes the time evolution of a density of an ensemble of systems. Its solution, given an initial density $f(t = 0) = f_0$, is

$$f(\cdot, t) = \exp(tL)f_0 = P(t)f_0, \tag{2}$$

where $P(t)$ is the Perron-Frobenius operator. It can be understood as the propagator of the probability density $f_0$. In its decomposed form for two systems $A$ and $B$, the operator is written as

$$f(x, t) = (P_A(t) \otimes P_B(t))\, f_0(x) = P(t)f_0(x). \tag{3}$$

##### MSM transition matrix decomposition

The Perron-Frobenius operator $P$ can be approximated by a Markov model. The Markov model formulation propagates probability densities between discrete states; therefore, the problem requires performing a Galerkin discretization of $P$ using a discrete basis set. This can be done by partitioning the phase space completely or into the metastable regions, say $\{A_1, ..., A_k\}$. The most common basis set are indicator functions on these regions,

$$\mathbb{1}_{A_i(x)} = \begin{cases} 1 & \text{if } x \in A_i \\ 0 & \text{else.} \end{cases} \tag{4}$$

The Galerkin discretization will yield a low rank approximations of the operator $P$. When using indicator functions, the output will usually be in the form of a discrete-time MSM [2]. Assume that the dynamics of interest can be separated into two independent disjoint regions in phase space. The Galerkin discretization of Eq. 2 in each region yields two MSMs,

$$f_A(t + \tau) = T_A f_A(t), \qquad f_B(t + \tau) = T_B f_B(t), \tag{5}$$

where $\tau$ is the lagtime; $f_A$ and $f_B$ are the probability vectors of the corresponding MSMs; and $T_A$, and $T_B$ are the corresponding transition probability matrices. In this case, the matrices $T_A$ and $T_B$ approximate Perron-Frobenius operators, so following Eq. 3, the solution of the whole systems is given by

$$f(t+\tau) = (T_A \otimes T_B)\, f_0(t), \tag{6}$$

with $f_0 = f_{0A} \otimes f_{0B}$. The individual transition probability matrices are linear maps given by $T_A : \mathbb{R}^k \to \mathbb{R}^k$ and $T_B : \mathbb{R}^{k'} \to \mathbb{R}^{k'}$, where $k$ and $k'$ are the number of discretized states in $A$ and $B$, respectively. The joint space $A \otimes B$ will then have $kk'$ discretized states that are defined analogously to Eq. 4 by indicator functions $\mathbb{1}_{(A\otimes B)_{(i,j)}(x)} = 1$ iff $x \in A_i \cap B_j$, so

$$T_A \otimes T_B : \mathbb{R}^{kk'} \to \mathbb{R}^{kk'}. \tag{7}$$

In particular, the product $T_A \otimes T_B$ is the Kronecker product [3]. An analogous expression can be derived for continuous-time MSMs (see below).

In summary, we need the operator $P$ to model the full system, and approximate it by the Kronecker product between the transition probability matrices of the MSMs of independent sub-systems.

#### Observable operator decomposition

To score dependency of subsystems in feature space, it is most natural to directly work with the operator that propagates these features. This operator is called the Koopman operator $K$; it propagates observable functions $f$,

$$Kf(x) = \mathbb{E}[f(\Phi(x))], \tag{8}$$

i.e., is described by the expectation value of the observable of a particular configuration, $x$, after the dynamics $\Phi$ has been applied. It is a infinite-dimensional linear operator [4]. It is particularly interesting for the current application because the variational approach for Markov processes (VAMP) and the related VAMP scores are derived from the Koopman operator [5].

As the Koopman formulation is a more general framework to deal with Markov processes, we only refer to the estimator of the Koopman operator which reads [5]

$$K = {C_{00}}^{-1/2} C_{0t} {C_{tt}}^{-1/2} \tag{9}$$

with time-lagged covariance matrix $C_{0t}$, "instantaneous" covariance matrices at times 0 and $t$ $C_{00}$ and $C_{tt}$, respectively. The lag time is $t$.

We define the common space of observables of two processes as a stacked vector $\Psi_{AB} = [\Psi_A, \Psi_A]$. For example, if $\Psi_A = (\psi_A^1, \psi_A^2, \ldots)$ and $\Psi_B = (\psi_B^1, \psi_B^2, \ldots)$ are the one-dimensional time series of features $\psi \in \mathbb{R}$ of two systems $A$ and $B$, the joint space would be spanned by $\Psi_{AB} = ((\psi_A^1, \psi_B^1), (\psi_A^2, \psi_B^2), \ldots)$. Note that this means that the separation of processes happens *a priori* by the choice of $\Psi_A, \Psi_B$, a situation which comes closest to applied modeling situations. Given the above definition of the full system observable $\Psi_{AB}$, the full system Koopman operator is the direct sum of Koopman sub-operators $K_A, K_B$, $K_{AB} = K_A \oplus K_B$ or, more generally,

$$K \begin{pmatrix} \Psi_1 \\ \Psi_2 \\ \vdots \\ \Psi_n \end{pmatrix} (x) = \begin{pmatrix} K_1 & 0 & \ldots & 0 \\ 0 & K_2 & \ddots & \vdots \\ \vdots & \ddots & \ddots & 0 \\ 0 & \ldots & 0 & K_n \end{pmatrix} \begin{pmatrix} \Psi_1 \\ \Psi_2 \\ \vdots \\ \Psi_n \end{pmatrix} (x) \tag{10}$$

as all off-diagonal blocks must vanish by definition and each subsystem operator only acts on the features of its space. Therefore, the decomposition can be written as the direct sum

$$K = \bigoplus_i K_i. \tag{11}$$

In particular, this means that the Koopman operator has the shape of a block diagonal matrix. It can be seen from the Koopman estimator (Eq. 9) that the above structure of the joint operator implies that independent processes are also uncorrelated.

### VAMP score decomposition of independent systems

The VAMP-$p$ score $R_p$ can be interpreted as the Schatten-$p$ norm $\|\cdot\|_p$ of the estimated Koopman operator to the $p$-th power [5], i.e.

$$R_p(K) = \|K\|_p^p. \tag{12}$$

This general form is valid for both MSMs as well as Koopman models, but note that the estimator for $K$ is different in these cases (see below). To simplify this expression, on the one hand, we can exploit the property of the Schatten-$p$ norm to be invariant under unitary transformations for unitarian matrices $U$ and $V$,

$$\|A\|_p = \|UAV\|_p. \tag{13}$$

On the other hand, we can write the Koopman operator in a singular value decomposition with its singular value diagonal matrix $\Lambda$ as $K = U\Lambda V$ such that, using Eq. 13, we find

$$\|K\|_p = \|\Lambda\|_p = \left(\sum_i \lambda_i^p\right)^{\frac{1}{p}} \tag{14}$$

with the real valued singular values of the Koopman matrix $\lambda_i$.

#### Sum space decomposition

Given a joint space that is spanned by the direct sum of subspaces, such as described with molecular observable vectors, and a decomposable Koopman operator $K_{AB} = K_A \oplus K_B$ of two systems $A$ and $B$, we can thus write

$$\begin{aligned} K_{AB} &= U_{AB}\Lambda_{AB}V_{AB} && (15)\\ &= (U_A \oplus U_B)(\Lambda_A \oplus \Lambda_B)(V_A \oplus V_B) && (16)\\ &= (U_A\Lambda_A V_A) \oplus (U_B\Lambda_B V_B) && (17)\\ &= K_A \oplus K_B && (18) \end{aligned}$$

and hence

$$\begin{aligned} \|K_{AB}\|_p^p &= \|\Lambda_{AB}\|_p^p && (19)\\ &= \|\Lambda_A \oplus \Lambda_B\|_p^p. && (20) \end{aligned}$$

Further, the singular values of the direct sum joint operator are the set of subsystem operator singular values. In detail, writing the $p$-th power of the $p$-Schatten norm of a real valued diagonal matrix (Eq. 14) reads

$$\left\|\begin{pmatrix} \lambda_{A,1} & 0 & \cdots & 0 \\ 0 & \ddots & & \vdots \\ \vdots & & \lambda_{B,1} & 0 \\ 0 & \cdots & 0 & \ddots \end{pmatrix}\right\|_p^p = \mathrm{Tr}\begin{pmatrix} \lambda_{A,1}^p & 0 & \cdots & 0 \\ 0 & \ddots & & \vdots \\ \vdots & & \lambda_{B,1}^p & 0 \\ 0 & \cdots & 0 & \ddots \end{pmatrix}$$

such that it follows that we can further simplify Eq. 20 to

$$\begin{aligned} &= \|\Lambda_A\|_p^p + \|\Lambda_B\|_p^p && (21)\\ &= \|K_A\|_p^p + \|K_B\|_p^p && (22) \end{aligned}$$

which is the VAMP-$p$ score of two independent systems in this particular basis. We can see that the decomposability depends on the block diagonal shape of the joint Koopman operator, which is also inherent to the covariance matrix itself. I.e., a decomposition of the covariance matrix would be possible in the same way, however its trace and Frobenius norm do not represent VAMP scores.

#### Product space decomposition

When operating in a joint space that is spanned by the tensor product, as shown above, the joint operator is formed by the Kronecker product $T_{AB} = T_A \otimes T_B$. However, the VAMP-score of a transition matrix $T$ is not directly computed from $T$ but from the associated Koopman operator. We first show that a decomposition of $T_{AB} = T_A \otimes T_B$ also implies a decomposition of $K_{AB}$ in the same way. We note that that the instantaneous correlation matrices are diagonal for MSMs. In the following, we make use of the transition matrix estimator $T = C_{00}^{-1} C_{0t}$ and the mixed product rule of Kronecker products.

$$K_{AB} = {}^{AB}C_{00}^{1/2} T_{AB} {}^{AB}C_{tt}^{-1/2} \tag{23}$$

$$= \left({}^{A}C_{00}^{1/2} \otimes {}^{B}C_{00}^{1/2}\right) (T_A \otimes T_B) \left({}^{A}C_{tt}^{-1/2} \otimes {}^{B}C_{tt}^{-1/2}\right) \tag{24}$$

$$= \left({}^{A}C_{00}^{1/2} T_A {}^{A}C_{tt}^{-1/2}\right) \otimes \left({}^{B}C_{00}^{1/2} T_B {}^{B}C_{tt}^{-1/2}\right) \tag{25}$$

$$= K_A \otimes K_B \tag{26}$$

Please note that this simple proof is only valid for indicator function basis sets such as for classical MSMs.

We can further make use of a simple rule that applies to the singular value decomposition of the Kronecker product. If the subsystem operators have $n$ and $m$ singular values $\lambda_{A,i} \in \mathbb{R}$ and $\lambda_{B,i} \in \mathbb{R}$, respectively, the singular values of its Kronecker product are $\{\lambda_{A,i} \cdot \lambda_{B,j} : 0 < i < n, 0 < j < m\}$. It thus follows that

$$\|K_{AB}\|_p^p = \sum_i \lambda_{AB,i}^p \tag{27}$$

$$= \sum_{i,j} (\lambda_{A,i} \cdot \lambda_{B,j})^p \tag{28}$$

$$= \sum_i \lambda_{A,i}^p \cdot \sum_j \lambda_{B,j}^p \tag{29}$$

$$= \|K_A\|_p^p \cdot \|K_B\|_p^p. \tag{30}$$

This is the decomposition for the VAMP-$p$ score of two independent systems in a product basis such as the one applied for MSM transition matrices.

#### Continuous-time MSM decomposition

Discretizations of the operator $P$ can also yield continuous-time MSMs [6, 7]. Analogously to the analysis done with discrete time MSMs, assume two independent regions in phase space that are discretized into two continuous-time MSMs. Their solution is

$$f_A(t) = \exp(tR_A) f_{0A}, \qquad f_B(t) = \exp(tR_B) f_{0B}, \tag{31}$$

where $R_A$ and $R_B$ are the transition rate matrices; $f_A$ and $f_B$ the probability densities in the corresponding regions; and $f_{0A}$ and $f_{0B}$ the initial conditions. The operator $P$ is approximated by the exponential functions, so the solution of the whole system is given by the tensor product of exponentials,

$$f(t) = \exp(t(R_A \oplus R_B)) f_0, \tag{32}$$

which yields a Kronecker sum $\oplus$ for the matrices in the exponent.

In summary, in order to approximate the operator $P$ of the full system, we need to either use the Kronecker sum on rate matrices of continuous-time MSMs, or the Kronecker product on transition probability matrices of discrete-time MSMs. In general, the full system Perron-Frobenius operator can be reassembled by using the tensor product on all the subsystems operators.

#### Weakly coupled systems

Practical situations – for example, an ion channel with quasi-independent subunits – might often involve weak coupling. The transition matrix $\widetilde{T}$ of a weakly coupled system can be expressed as a perturbation of the transition matrix $T$ of the non-coupled system,

$$\widetilde{T}^{\mathrm{T}} = (1 - \epsilon) T^{\mathrm{T}} + \epsilon P^{\mathrm{T}}, \quad \epsilon \in [0, 1] \tag{33}$$

where $P$ is another Markov transition matrix defined on the same state space as $T$, and $\epsilon \ll 1$ corresponds to small perturbations/weak coupling. Note this definition enforces the required MSM condition that columns sum to one.

As the eigenvalues of $\widetilde{T}$ are continuous functions of $\epsilon$, the eigenvalues of the coupled system will be arbitrarily close to those of the uncoupled one as $\epsilon \to 0$. Further analysis on the convergence speed of the eigenvalues as $\epsilon \to 0$ is system dependent and not easy to assess in general. However, upper error bounds for the stationary distribution error exist and can be assessed in multiple ways [8, 9]. We focus on one formulation framed in terms of mean first passage times $m_{ij}$, since it provides physical intuition on the sensitivity of the MSM [8, 10]. Assume $T$ and $P$ define finite, irreducible and homogeneous MSMs, as the MSMs of interest within the scope of this work, then

$$\|\pi - \tilde{\pi}\|_\infty \leq \frac{1}{2} \max_j \left[ \frac{\max_{i \neq j} m_{ij}}{m_{jj}} \right] \|(T - \widetilde{T})^\mathrm{T}\|_\infty, \tag{34}$$

where $\pi$ denotes the stationary distribution; the tilde denotes quantities of the perturbed system; the $\infty$-norm is the maximum absolute row sum, and $m_{jj}$ is the mean return time of state $j$, i.e. the time to return to $j$ for the first time, starting from $j$.

In terms of our application, if the coupling is sufficiently weak ($\epsilon \ll 1$), the eigenvalues of the uncoupled system will be close to those of the weakly coupled system, providing a good approximation of the implied timescales. Furthermore, an upper bound for the stationary distribution error can be easily calculated using software like PyEMMA [11]. The bound is very effective for MSMs consisting of a dominant central state with strong connections to and from all other states [8].

### Toy models

#### Scaling behavior: 2 uncoupled 3 state sub-systems

A system consisting of $n$ independent sub-systems with 3 states each was set up to exemplify scaling with number of sub-systems. The transition matrix of each sub-system is given by

$$T_i = \begin{pmatrix} 1-p & p/2 & p/2 \\ p/2 & 1-p & p/2 \\ p/2 & p/2 & 1-p \end{pmatrix} \tag{35}$$

with the probability $p = 0.1$ to stay in a particular state. The full system is described with a Kronecker product $T_\text{full} = \bigotimes_i^n T_i$. Markov chains of length $N$ are sampled from this transition matrix using PyEMMA / msmtools [11] until the desired set of states is connected. To quantify the confidence, 30 trial runs are conducted for each number of sub-systems.

#### Approximation quality: 2 weakly coupled 2 state sub-systems

In the following, we utilize a system comprised of 2 sub-systems with 2 states each in order to exemplify the IMD framework. We further analyze its behavior with regard to limited sampling and weak couplings. The toy model consists of two sub-systems with transition matrices $T_1, T_2$ that each have a probability to transition to another state of $\epsilon = 0.1$.

These sub-systems are coupled in a tunable fashion. A parameter $\lambda$ is introduced which results in two independent sub-systems for $\lambda = 0$ and weakly coupled sub-systems for $\lambda > 0$. The full system is represented by a reversible transition matrix $T$ for any given $\lambda \in ]0, \epsilon(1-\epsilon)[$. The transition matrix of the applied toy model can explicitly be written as

$$T = \begin{pmatrix} (1-\epsilon)^2 - \lambda & \epsilon(1-\epsilon) - \lambda & \epsilon(1-\epsilon) + \lambda & \epsilon^2 + \lambda \\ \epsilon(1-\epsilon) - \lambda & (1-\epsilon)^2 - \lambda & \epsilon^2 + \lambda & \epsilon(1-\epsilon) + \lambda \\ \epsilon(1-\epsilon) + \lambda & \epsilon^2 + \lambda & (1-\epsilon)^2 - \lambda & \epsilon(1-\epsilon) - \lambda \\ \epsilon^2 + \lambda & \epsilon(1-\epsilon) + \lambda & \epsilon(1-\epsilon) - \lambda & (1-\epsilon)^2 - \lambda \end{pmatrix} \tag{36}$$

$$\stackrel{\lambda=0}{=} T_1 \otimes T_2. \tag{37}$$

In the un-coupled case, $T$ reduces to the Kronecker product of the two sub-system transition matrices. The sub-system transition matrices are given by

$$T_1, T_2 = \begin{pmatrix} 1-\epsilon & \epsilon \\ \epsilon & 1-\epsilon \end{pmatrix}. \tag{38}$$

We sample discrete trajectories from $T$, de-compose into sub-system trajectories and estimate the models presented in Fig. 1.

As expected, all properties of the Markov model can be easily retained in the uncoupled case (Fig. 1). Stronger coupling yields less accurate results; especially transition probabilities are over or underestimated (Fig. 1a) while the error on the implied timescales is comparably small, possibly yielding underestimated implied timescales (Fig. 1b). We note that the stationary probabilities are not affected by the coupling, i.e. that $p_1 \cdot p_2 = p_{1,2}$ holds in any case (Fig. 1c). We find that indeed, the *dependency* $d$ in both its forms, trace and Frobenius norms, is a fast converging and significant indicator for the approximation quality (Fig. 1d).

Due to its small size, this particular example is not suitable to demonstrate that convergence is reached faster with the decomposed model.

### Dimer model

The following model system serves the purpose to demonstrate that the presented dependency scores can bisect coupled from weakly coupled systems. Our example models a dimer of protein channels. Each of those channels resembles a Hodgkin-Huxley potassium channel but possesses an additional deactivation switch. This switch alters the dynamics completely, i.e. upon activation each gate will close or stay closed with a high probability. The deactivation switch is a Markov process itself and switches state with a probability of $p_{\text{switch}} = 0.01$. Thus, each channel has strongly coupled sub-units and cannot be described by individual gate MSMs as in the previous example.

Our test system consists of two such channels. They possess some weak cooperativity which we model by a slight shift in gate opening probability if both deactivation switches are disabled at the same time.

In the following, we define a block matrix that describes the whole system dynamics. For the sake of simplicity, we present it in multiple layers. The highest layer describing the full system is given by

$$T_{\text{dimer}} = \begin{pmatrix} T\begin{pmatrix} S1:0\to 0 \\ S2:0\to 0 \end{pmatrix} & T\begin{pmatrix} S1:0\to 0 \\ S2:0\to 1 \end{pmatrix} & T\begin{pmatrix} S1:0\to 1 \\ S2:0\to 0 \end{pmatrix} & T\begin{pmatrix} S1:0\to 1 \\ S2:0\to 1 \end{pmatrix} \\ T\begin{pmatrix} S1:0\to 0 \\ S2:1\to 0 \end{pmatrix} & T\begin{pmatrix} S1:0\to 0 \\ S2:1\to 1 \end{pmatrix} & T\begin{pmatrix} S1:0\to 1 \\ S2:1\to 0 \end{pmatrix} & T\begin{pmatrix} S1:0\to 1 \\ S2:1\to 1 \end{pmatrix} \\ T\begin{pmatrix} S1:1\to 0 \\ S2:0\to 0 \end{pmatrix} & T\begin{pmatrix} S1:1\to 0 \\ S2:0\to 1 \end{pmatrix} & T\begin{pmatrix} S1:1\to 1 \\ S2:0\to 0 \end{pmatrix} & T\begin{pmatrix} S1:1\to 1 \\ S2:0\to 1 \end{pmatrix} \\ T\begin{pmatrix} S1:1\to 0 \\ S2:1\to 0 \end{pmatrix} & T\begin{pmatrix} S1:1\to 0 \\ S2:1\to 1 \end{pmatrix} & T\begin{pmatrix} S1:1\to 1 \\ S2:1\to 0 \end{pmatrix} & T\begin{pmatrix} S1:1\to 1 \\ S2:1\to 1 \end{pmatrix} \end{pmatrix}. \quad (39)$$

Its block elements depend on deactivation switches of the individual channels, $S1$ and $S2$. On the next layer, for each transition pair of the deactivation switches,

$$T\begin{pmatrix} S1:i\to j \\ S2:n\to m \end{pmatrix} = \begin{cases} T_c \otimes T_c & \text{if } n=m=i=j=0 \\ T(S1:i\to j)\otimes T(S2:n\to m) & \text{else.} \end{cases} \quad (40)$$

This implements the coupling between channels by selecting different transition probabilities if both deactivation gates are inactive at the same time. The next layer describes individual channel transition probabilities (rescaled such that the full system transition matrix has row-sum 1):

$$T(S:i\to j) = \begin{cases} p_{\text{switch}} \cdot \mathbb{1}_{16} & n \neq m \text{ switching switch} \\ (1-p_{\text{switch}}) \cdot T_{\text{HH}} & n=m=0 \text{ inactive switch} \\ (1-p_{\text{switch}}) \cdot T_{\text{inactive}}(\lambda) & n=m=1 \text{ active switch} \end{cases} \quad (41)$$

with the Hodgkin-Huxley transition matrix $T_{\text{HH}} = T_{hh} \otimes T_{hh} \otimes T_{hh} \otimes T_{hh}$ with individual gate transition matrices $T_{hh}$ that describe gate opening and closing in the native state. Further, a transition matrix describing gate dynamics if the deactivation switch is active is given. For fully activated coupling between gates and deactivation switch, it reads $\tilde{T}_{\text{inactive}} = T_o \otimes T_o \otimes T_o \otimes T_o$ with $T_o$ being the single gate transition matrices for that case. In order to control the intensity of the gate-deactivation switch coupling, we use a linear mixture parameter $\lambda$,

$$T_{\text{inactive}}(\lambda) = \lambda \tilde{T}_{\text{inactive}} + (1-\lambda) T_{\text{HH}}, \quad (42)$$

i.e., the coupling can gradually be turned off by adjusting $\lambda \in ]0,1[$, and the case $\lambda = 0$ leaves the deactivation switch with no effect on the gate dynamics.

Finally, on the last layer, the single gate matrices are given by

$$T_{hh} = \begin{pmatrix} 0.9483 & 0.0517 \\ 0.0055 & 0.9945 \end{pmatrix} \qquad \text{unperturbed} \tag{43}$$

$$T_o = \begin{pmatrix} 0.9483 & 0.0517 \\ 0.95 & .05 \end{pmatrix} \qquad \text{active deactivation switch} \tag{44}$$

$$T_c = \begin{pmatrix} 0.8 & 0.2 \\ 0.0055 & 0.9945 \end{pmatrix} \qquad \text{both deactivation switches inactive} \tag{45}$$

The Markov chain is sampled from the transition matrix $T_{\mathrm{dimer}}$ using PyEMMA / msmtools [11] with a time step of 20 steps for 1 million time steps. The code used to generate and analyze the example can be found in our GitHub repository.

179 # Supplementary figures

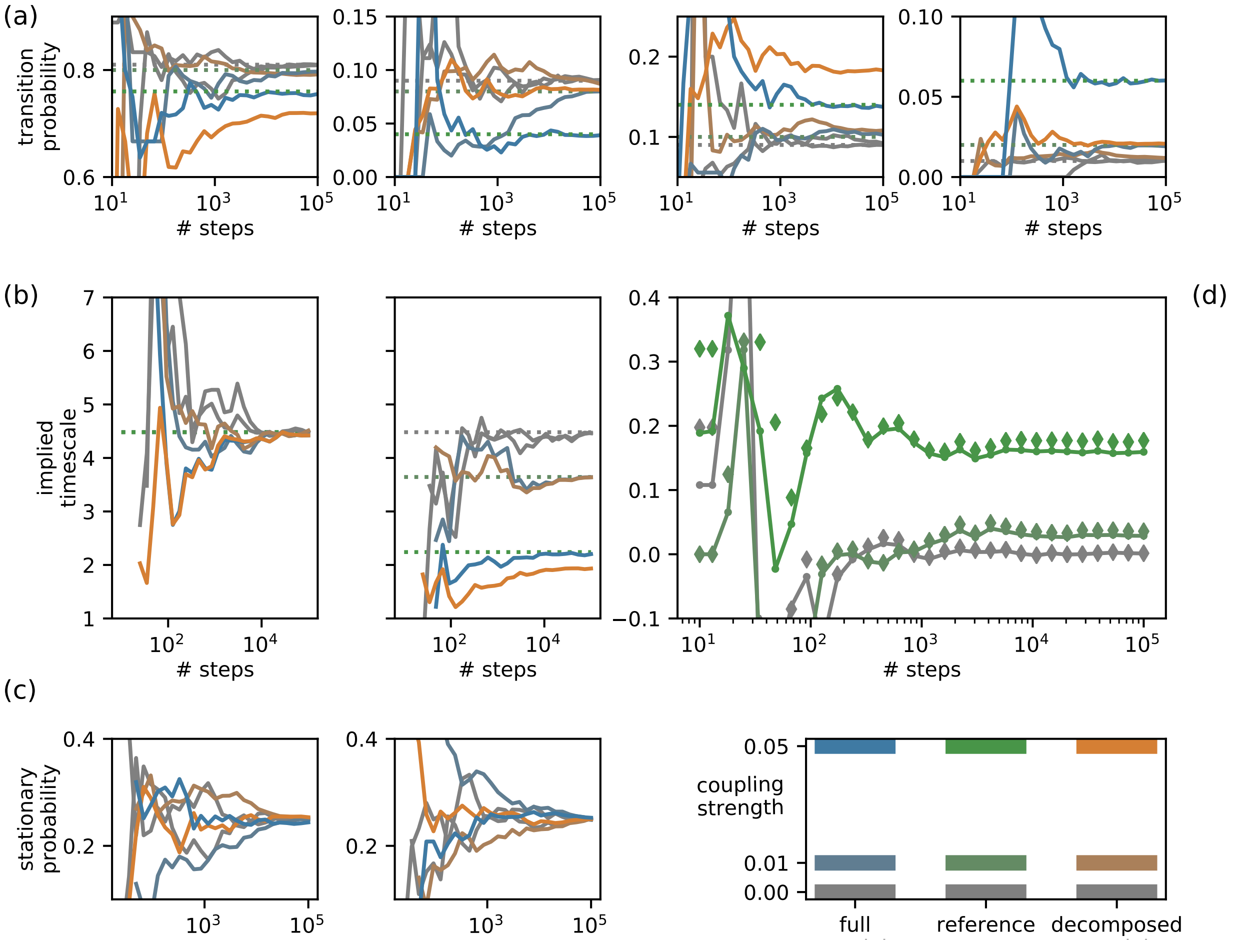


**Figure S1:** Analysis of error from weak couplings and limited sampling. MSM properties of full-system and decomposed estimates are shown as functions of sampling (x-axis) and coupling (color code). (a) first row of transition probability matrix, (b) two highest implied timescales. (c) Stationary probabilities (shown for two example states). (d) *dependency* $d$ as difference in trace norms (line) and Frobenius norms (diamonds).

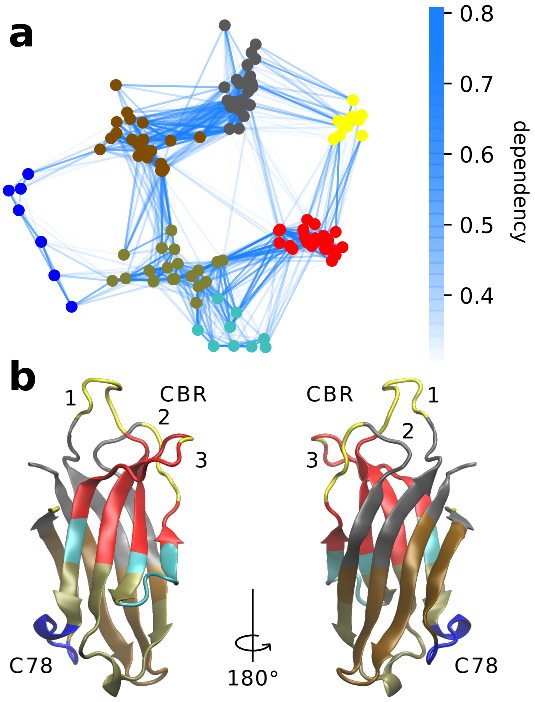


**Figure S2:** Dependency-network between residues of Syt-1 C2A depicted using a standard graph layout (Fruchterman-Rheingold algorithm). a: VAMP-1 normalized dependency network. Edge weights are indicated by colorbar. Nodes are colored according to an unsupervised classification by the $k$-means algorithm ($k = 7$). b: Visualization of protein structure with color coded segments from our VAMP-1 analysis, i.e., same color code as in panel a.
